# Meta-analysis of Cytometry Data Reveals Racial Differences in Immune Cells

**DOI:** 10.1101/130948

**Authors:** Zicheng Hu, Chethan Jujjavarapu, Jacob J. Hughey, Sandra Andorf, Hao-Chih Lee, Pier Federico Gherardini, Matthew H. Spitzer, Patrick Dunn, Cristel G. Thomas, John Campbell, Jeff Wiser, Brian A. Kidd, Joel T. Dudley, Garry P. Nolan, Sanchita Bhattacharya, Atul J. Butte

## Abstract

While meta-analysis has demonstrated increased statistical power and more robust estimations in studies, the application of this commonly accepted methodology to cytometry data has been challenging. Different cytometry studies often involve diverse sets of markers. Moreover, the detected values of the same marker are inconsistent between studies due to different experimental designs and cytometer configurations. As a result, the cell subsets identified by existing auto-gating methods cannot be directly compared across studies. We developed MetaCyto for automated meta-analysis of both flow and mass cytometry (CyTOF) data. By combining clustering methods with a silhouette scanning method, MetaCyto is able to identify commonly labeled cell subsets across studies, thus enabling meta-analysis. Applying MetaCyto across a set of 10 heterogeneous cytometry studies totaling 2926 samples enabled us to identify multiple cell populations exhibiting differences in abundance between White and Asian adults. Software is released to the public through GitHub (github.com/hzc363/MetaCyto).

## Main text

Meta-analysis of existing data across different studies offers multiple benefits. The aggregated data allows researchers to test hypotheses with increased statistical power. The involvement of multiple independent studies increases the robustness of conclusions drawn. In addition, the complexity of aggregated data allows researchers to test or generate new hypotheses. These benefits have been shown by many studies in areas such as genomics, cancer biology and clinical research and have led to important new biomedical findings ^1–4^. For example, one study showed the correlation between neoantigen abundance in tumors and patient survival by performing meta-analysis of RNA sequencing data from The Cancer Genome Atlas (TCGA)^5^. In another study, meta-analysis of genome-wide association studies identified novel loci that affect the risk of type 1 diabetes^6^.

With the recent advances in high-throughput cytometry technologies the immune system can be characterized at the single cell level with up to 45 parameters, minimizing the technical limitations and allowing capture of more valuable information from immunology studies^7–9^. Open science initiatives have led to more of this type of research data accessible, and the availability of shared cytometry data, including data from flow cytometry and mass cytometry (CyTOF), is growing exponentially. Notably, the ImmPort database (www.immport.org), a repository for immunology-related research and clinical trials, provides numerous studies with thousands of cytometry datasets^10^. However, meta-analysis of cytometry datasets remains particularly challenging. Different studies use diverse sets of protein markers and fluorophore/isotope combinations. The detected values of the same marker are inconsistent between studies because of different cytometer configurations or operators. In addition, the high dimensionality of cytometry data, especially CyTOF data, makes manual gating based meta-analysis difficult and time-consuming.

Multiple computational methods have been proposed to automate the analysis of cytometry data, such as FlowSOM^11^, FlowMeans^12^ and CITRUS^13^. Although they are effective in analyzing data from a single study^14,15^, several limitations have prevented their use in metaanalysis. First, the results of these methods cannot be directly compared across studies. The cell subsets identified by these methods are usually labeled with anonymous identifiers with no cell-specific annotation, making it impossible to identify common cell populations across different studies. In addition, many clustering methods are sensitive to parameter choices. For example, FlowSOM, FlowMeans and SPADE^16^ require users to pre-specify the number of clusters. As a result, extensive parameter tuning and manual inspection are required for every cytometry dataset. In meta-analysis where large numbers of input datasets could be involved, these manual selected choices become a major technical burden.

In this study, we developed MetaCyto to enable automated metaanalysis of cytometry datasets, including data from both conventional flow and CyTOF cytometry data. Using novel computational approaches, MetaCyto accurately identifies common cell populations across studies without parameter tuning requirements. It then applies hierarchical models to robustly estimate the effects of factors of interest, such as age, race or vaccination, on the cell populations using data across all input studies.

To test the utility of MetaCyto, we performed a joint analysis of 10 human immunology cytometry datasets contributed by four different institutions^17–20^. Altogether, this analysis spanned 2926 peripheral blood mononuclear cells (PBMC) or whole blood samples from 984 healthy subjects, which were acquired using either flow cytometry or CyTOF with a diverse set of markers. Among these 984 subjects, over 90 percent were White or Asian. While it is well known that characteristics of multiple immune system-related diseases, such as HIV^21^, systemic lupus erythematosus^22^ and hepatitis C^23^, vary between the two racial groups, the heterogeneity of the immune system among the human population has made studying these differences difficult^24,25^. We hypothesized that a meta-analysis approach could lead to a better understanding of racial differences in the immune system. Using MetaCyto, we not only confirmed a known racial difference, but also identified new cell types whose frequencies differ between White and Asian.

## RESULTS

### MetaCyto identifies common cell subsets across studies

Our meta-analysis of cytometry data follows four steps: data aggregation, data pre-processing, identification of common cell subsets across studies, and statistical analysis (**Fig. 1a**). The third step, identification of common cell subsets across studies, has been one of the main technical challenges preventing automated meta-analysis. Therefore, while all four steps are automated and covered in the MetaCyto software system and documented in the online methods, here we primarily focus on describing our identification and relating of common cell subsets across studies.

**Figure 1:**
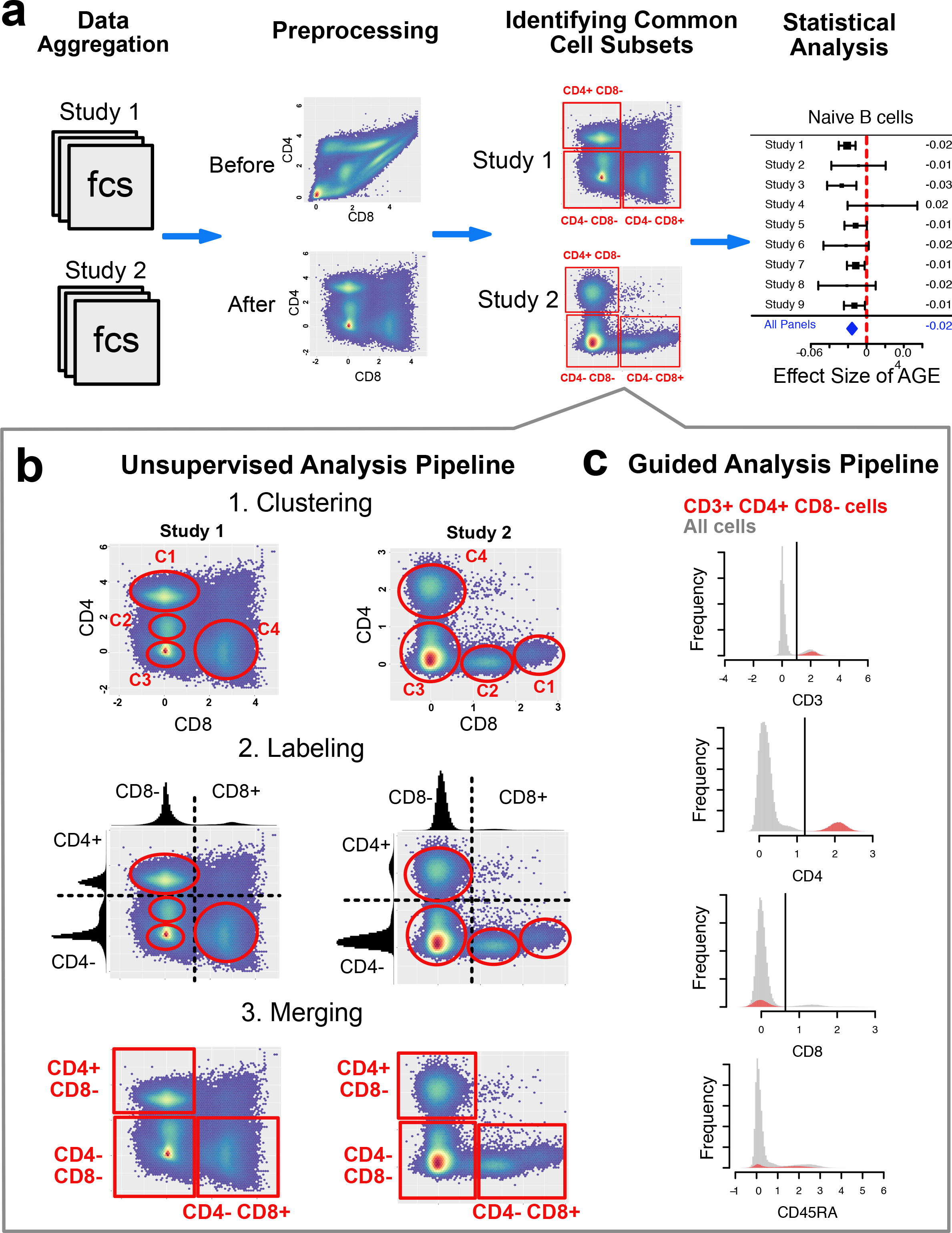
MetaCyto identifies and labels common cell subsets in cytometry data across studies. (**a**) Schematic illustration of the 4 steps MetaCyto uses to perform meta-analysis of cytometry data. (**b**) Schematic illustration of the unsupervised analysis pipeline in MetaCyto. Top: Cytometry data from different studies are first clustered using a clustering method, such as FlowSOM. Middle: Each marker is bisected into positive and negative regions using the silhouette scanning method. Each identified cluster is labeled based on their position relative to this threshold. Bottom: Clusters with the same label are merged together into rectangles or hyper-rectangles. (**c**) An example illustrating the guided analysis pipeline in MetaCyto. Each marker in the data is bisected into positive and negative regions using the silhouette scanning method. The CD3+ CD4+ CD8− cluster corresponds to cells that fall into CD3+ region, CD4+ region and CD8− region at the same time. Red histograms show the distribution of markers in CD3+ CD4+ CD8− subset. Grey histograms show the distribution of markers of all cells.

MetaCyto employs two automated pipelines, unsupervised analysis and guided analysis, to identify common cell subsets across studies. The unsupervised analysis pipeline identifies cell subsets in a fully automated way. Cytometry data in each study is first clustered using an existing clustering method (Fig. 1b **Top**). FlowSOM^11^ was implemented as the default clustering method due to its speed and performance. However, any other clustering method, such as hierarchical clustering or FlowMeans, could be substituted as well. At this stage, clusters are labeled with non-informative labels, such as C1, C2, C3, which cannot be related across studies. For example, C1 in study 1 and C1 in study 2 represent entirely different cell populations.

A threshold is then chosen to bisect the distribution of each marker into positive and negative regions, needed to label each cluster in a biologically meaningful way (Fig. 1b **Middle**). The selection of a threshold is easy when a clear bi-modal distribution is present, but becomes challenging in other cases. We implemented a Silhouette scanning method, which bisects each marker at the threshold maximizing the average silhouette, a widely used way of describing the quality of clusters^26^. We compared Silhouette scanning against 8 other bisection methods and found it to be superior (**Supplementary Fig. 1**).

Clusters are then labeled for each of the markers based on the following two rules: first, if the marker levels of 95% of cells in the cluster are above or below the threshold, the cluster will be labeled as positive or negative for the marker, respectively. Otherwise, the cluster will not be labeled for the marker. For example in Fig 1b, both C2 and C1 in study 2 will be labeled as CD8+ CD4−; second, if a marker is positive or negative in 95% of all cells, the marker is not used to label any clusters. For example, CD45, which is expressed by all immune cells, will not be used to label any cell clusters in the blood. The two rules are used to reduce redundancy and ensure that only the informative markers are used for labeling.

Next, clusters with the same labels are merged into a square shaped cluster (Fig. 1b **Bottom**). In cytometry data with higher dimensions, clusters are hyper-rectangles. Following this stage, common cell subsets across studies can be rigorously identified and annotated. For example, the CD4− CD8+ clusters in both study 1 and study 2 correspond to CD8+ T cells.

The guided analysis pipeline identifies cell subsets using pre-defined cell definitions, thus allowing for the search of specific cell subsets defined by immunologists. After bisecting each marker into positive and negative regions, cells fulfilling the pre-defined cell definitions are identified. For example, the CD3+ CD4+ CD8− (CD4+ T cells) cell subset corresponds to the cells that fall into the CD3+ region, CD4+ region and CD8- region concurrently (Fig. 1c). Notice that both CD45RA+ and CD45RA- populations are included in the cell subset, because the cell definition does not specify the requirement for CD45RA expression. However, researchers could easily alter the cell definition to CD3+ CD4+ CD8- CD45RA+ to find the CD45RA+ cell subset.

### Evaluating the guided analysis pipeline

A successful meta-analysis of cytometry data requires cell populations to be identified accurately from each study. To evaluate if the guided analysis pipeline of MetaCyto can accurately identify cell subsets from a single study, we downloaded a set of PBMC cytometry data (SDY478) from ImmPort, with which the original authors identified 88 cell types. Correspondingly, we specified the 88 cell definitions (**Supplementary Table 2**) based on the author’s gating strategy and identified these cell subsets for each cytometry sample using the guided analysis pipeline in MetaCyto. We compared the proportions of all cell subsets estimated by MetaCyto with the original manual gating results and found that MetaCyto estimations are highly consistent with the manual gating result (Fig. 2a-c). We compared our estimations to two existing methods, flowDensity^27^ and ACDC^28^, which can also identify pre-defined cell populations. Our results suggest that MetaCyto’s quantification of both major and rare populations were more accurate than FlowDensity’s (Fig. 2d,e). Although ACDC and MetaCyto results had the same correlation with manual gating, ACDC tended to overestimate the cell abundance (**Supplementary Fig. 2a,b**). In addition, a relatively shorter computational time of MetaCyto (around 3 minutes) compare to ACDC (over 2 hours) makes it advantageous in analyzing a large number of datasets.

**Figure 2:**
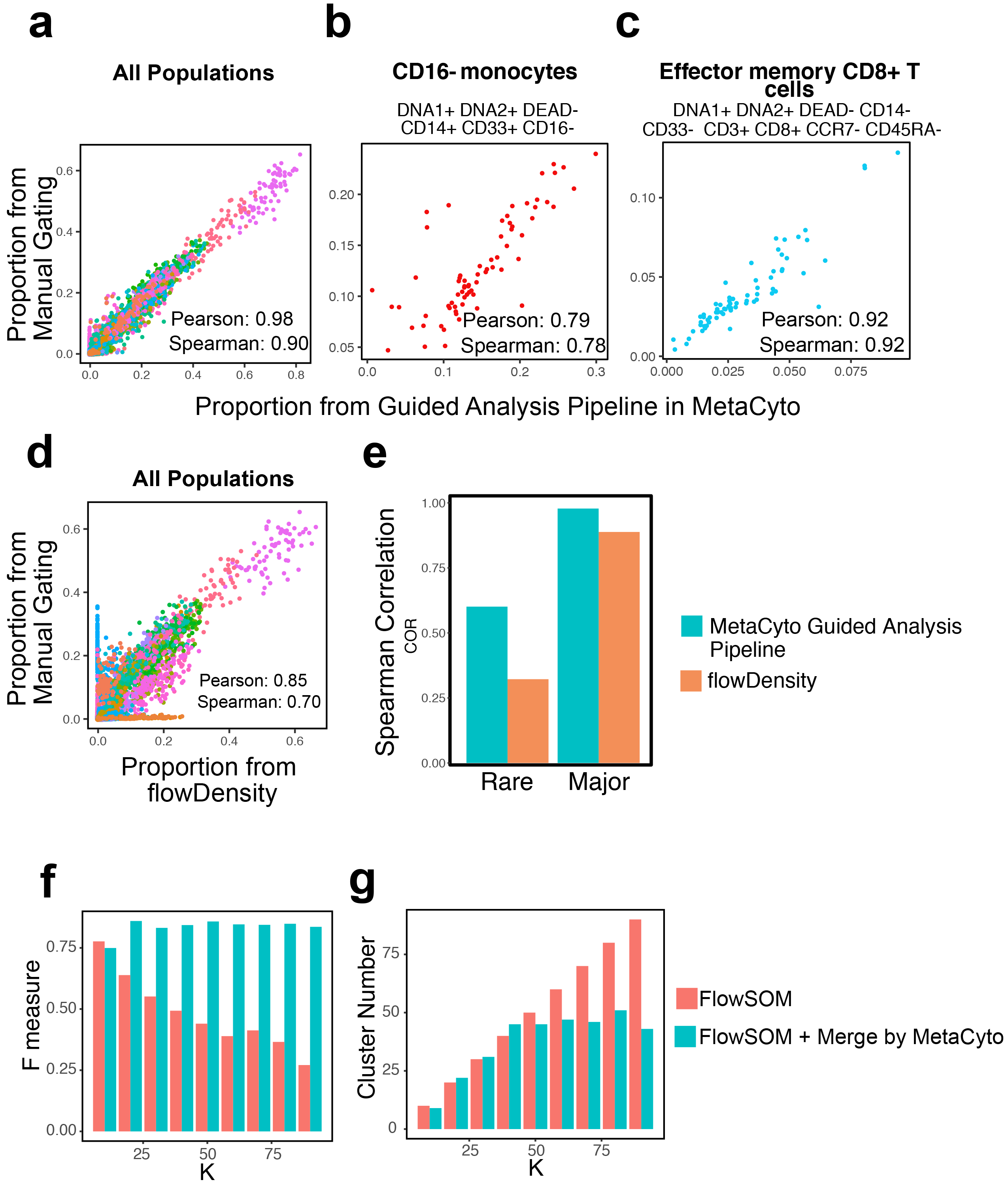
Both guided and unsupervised analysis pipelines in MetaCyto accurately identify cell populations. (**a-c**) Scatter plots showing the comparison between proportions of cell types estimated by the guided analysis pipeline in MetaCyto and proportions provided by the authors of SDY478. All cell populations (**a**), CD16− monocytes (**b**), and effector memory CD8+ T cells (**c**) are included in the plots. Each dot represents the proportion of a cell type in a sample. Each color represents a cell type. (**d**) Scatter plots showing the comparison between flowDensity and manual gating. All cell populations are included. (**e**) The 88 cell types are broken down into rare and major populations based on their mean proportion in the manual gating results. The cell types whose mean proportions are less than 2 percent are defined as rare population, the rest of the cell types are defined as major populations. Spearman correlation between MetaCyto or flowDensity’s results and manual gating results are calculated to measure the performance. (**f,g**) FlowSOM is used to cluster the West Niles Virus dataset (FlowCAP WNV) with K ranging from 10 to 90 with or without the merge step in MetaCyto unsupervised analysis pipeline. F measure (**f**) and the number of clusters (**g**) are shown in the bar plots. See also **Supplementary Fig. 2**.

### Evaluating the unsupervised analysis pipeline

We then tested the performance of the unsupervised analysis pipeline of MetaCyto. In the unsupervised analysis pipeline, cell clusters are first identified by an existing clustering algorithm. The subsets are then labeled using informative markers and are merged into hyper-rectangle clusters based on the labeling result (Fig. 1b). To learn how such merge affects the quality of clusters, we evaluated the results of two clustering algorithms, FlowSOM^11^ and FlowMeans^12^, with and without the merging step. Multiple studies have been conducted to evaluate the performance of existing clustering method for cytometry data^14,15^. The most recent (Weber et al.^15^) compared 15 clustering methods and found FlowSOM generally outperformed other methods after manual tuning.

We downloaded an evaluation dataset, West Nile virus dataset (FlowCAP WNV), used by Weber et al^15^ and applied FlowSOM. The clustering result is then labeled and merged. Since FlowSOM requires a pre-specified cluster number (K), we did multiple runs with K ranging from 10 to 90. F-measure is used to evaluate the quality of the clusters. We found that the quality of clusters is comparable before and after merging when K equals to 10. However, the performance of FlowSOM drops when K increases. The subsequent merging step prevented FlowSOM performance to deteriorate (Fig. 2f). We then looked at the total number of clusters identified before and after merging. As expected, FlowSOM identified the same number of clusters specified by K. However, when running the merging step after FlowSOM, the total number of clusters no longer increases after a certain point (Fig. 2g). The same results were obtained with FlowMeans^12^ (**Supplementary Fig. 2c,d**).

To see if such benefit of the merging step only exists in datasets where the intrinsic number of cell subsets is small, we applied the same methodology in the normal donor (ND) dataset from FlowCAP competition^14^, where more cell subsets can be identified. Consistent with results from WNV dataset, the merging step is able to prevent the over-partitioning in the ND dataset as well (**Supplementary Fig. 2e-f**).

The results suggest that MetaCyto is able to merge small clusters in a biologically meaningful way, preventing over-partitioning of the cell subsets, thus allowing the clustering analysis to be performed without tuning any parameters.

### Meta-analysis using MetaCyto confirms previous findings

After evaluating the performance of MetaCyto in analyzing cytometry data from single studies, we next tested the ability of MetaCyto in yielding consistent results from combining multiple studies. We applied MetaCyto to identify cell types whose frequencies are different between age, gender and race groups. We downloaded 10 studies from ImmPort containing cytometry data. These 10 studies had been contributed from four different institutions, where 86 panels containing 74 different markers were used (Fig. 3 and **Supplementary Table 3**). Altogether, the datasets contain 2926 whole blood or PBMC samples from 984 healthy subjects and were acquired using either flow cytometry or CyTOF. The subjects are proportionately distributed by gender, with slightly more female than male. The age span range from 19 to 90 years. The subjects come from five different defined racial groups. Among them, over 90% were White or Asian (**Supplementary Fig. 3**).

**Figure 3:**
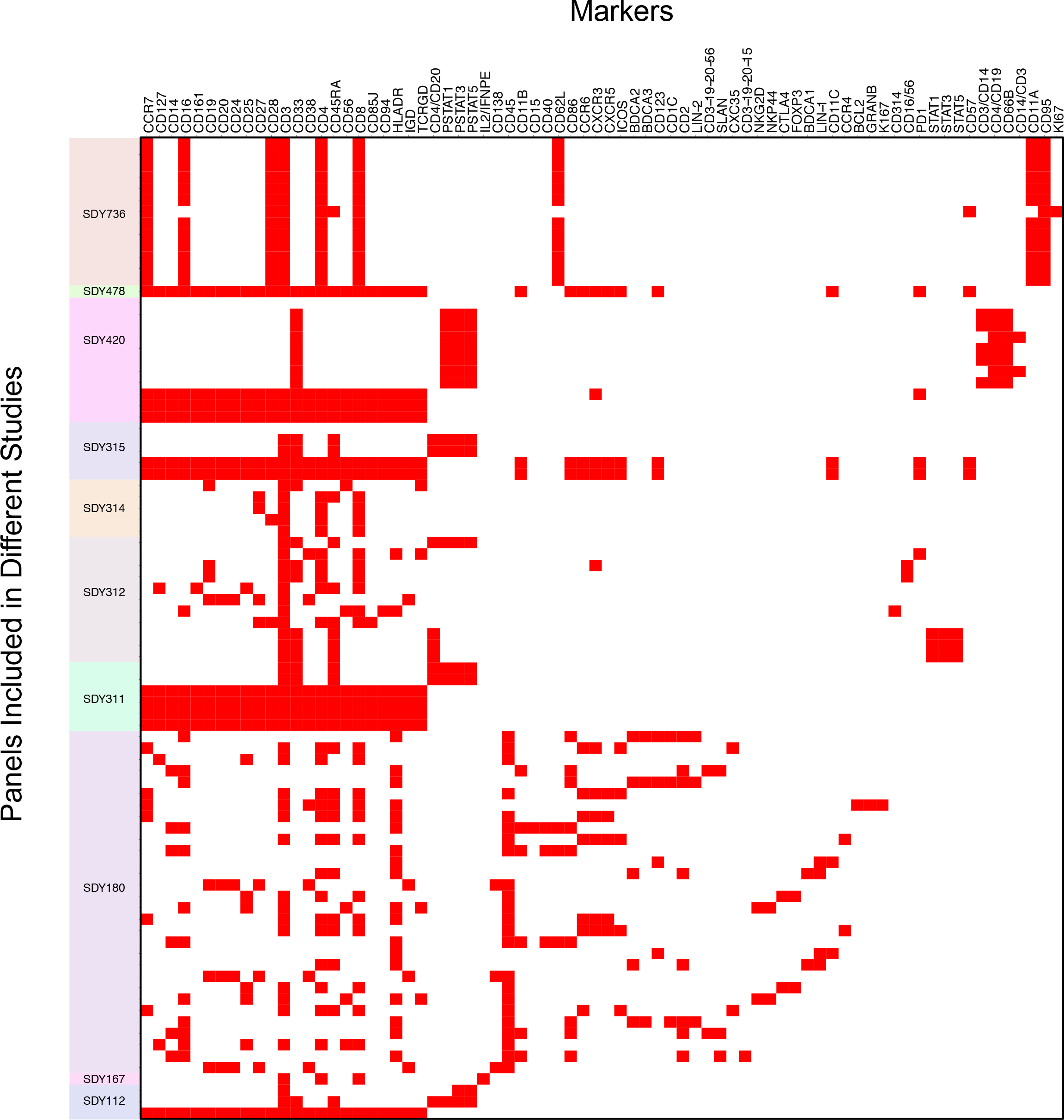
Data from 10 human immunology studies includes highly heterogeneous cytometry panels. Eighty-six panels with diverse sets of markers were used in these 10 studies, with the panels represented vertically. The specific markers used are represented horizontally. A red square in each grid element indicates that particular marker was used in a panel.

We used both unsupervised and guided MetaCyto analysis pipelines in parallel to identify cell subsets. For the latter, we used 23 cell type definitions from the Human ImmunoPhenotyping Consortium (HIPC)^29^, ranging from effector memory T cells to monocytes (**Supplementary Table 4**). We then estimated the effect size of age, gender and race on the cell type proportions using hierarchical statistical models.

To see if the results from MetaCyto were consistent across studies, we performed leave-one-out analysis ten times. Each time, we left one of the 10 studies out and estimated the effect sizes of age, gender and race using the rest nine studies. We found that the leave-one-out analysis agree well with the full meta-analysis (correlation ranges from 0.76 to 1, **Supplementary Table 5**).

We then validated our results using the effect sizes of age and gender, previously well characterized in other studies^20,30^. We tested whether results obtained with MetaCyto could replicate results from a previous independent study (Carr et al.^30^). Among the 23 cell types identified by the guided analysis pipeline, 14 overlapped with the cell types included in the Carr study. We compared the effect size of age and gender on the proportion of these 14 cell types, between MetaCyto on the 10 studies, and the independent results from Carr, et al. We found that results agree well with each other on both the effect size of age (r = 0.69, p = 0.006, Fig. 4a) and gender (r = 0.71, p = 0.004, Fig. 4b).

**Figure 4:**
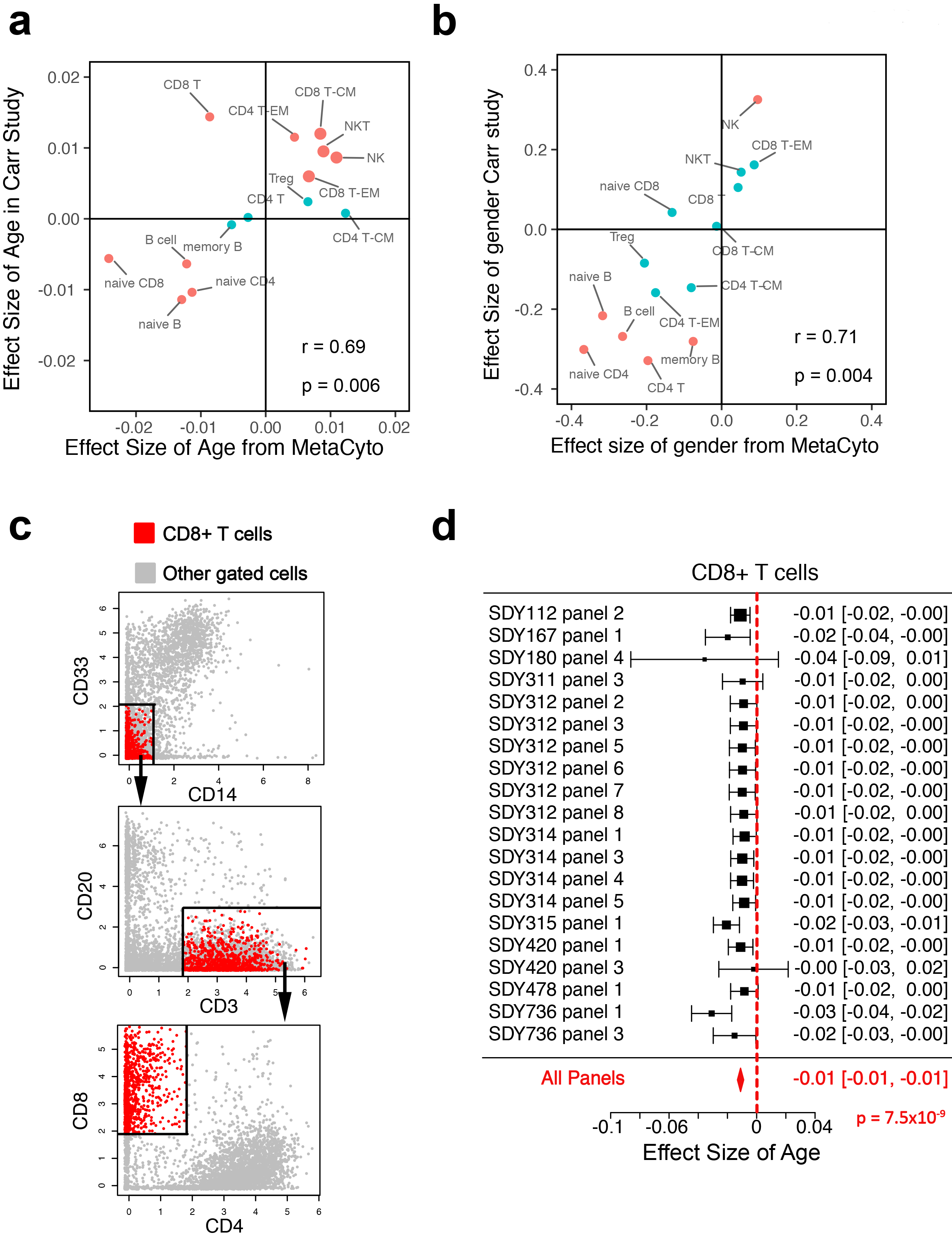
Meta-analysis using MetaCyto provides consistent results between cytometry panels and confirms previous findings. (**a-b**) Comparison between the effect sizes of age (**a**) and gender (**b**) estimated by MetaCyto using all 86 panels, against the effect sizes estimated using the data from Carr, et al. Red dots represent significant findings by MetaCyto. (**c**) 2D plots visualizing the CD8+ T cells identified by MetaCyto. Red dots represent the cells identified by MetaCyto. Grey dots represent other cells in the gate. Data from SDY420 are shown as an example. (**d**) A forest plot showing the effect size of race (Asian compared to White) on the proportion of CD8+ T cells in the blood. The effect sizes were estimated within each panel first, and are combined using a random effect model. In **a** and **b**, r represents the Pearson correlation; p represents the p value of r not equal to 0. In **d**, p was calculated using a random effect model.

The only discrepancy between our analysis and Carr study was the effect of age on CD8+ T cells (Fig. 4a). Our result showed that the proportion of CD8+ T cells significantly decreases with age, while Carr study reported an increase with age. We visually inspected MetaCyto’s auto-gating, and ruled out such disagreement being caused by gating errors in our study (Fig. 4c). The forest plot showed that our finding was consistent across cytometry panels (Fig. 4d). In the literature, one study found that CD8+ T cell proportion decrease with age^31^ while another study found no association between CD8+ T cells and age^32^. These discrepancies suggest that the effect of age on CD8+ T cells is highly variable and environment specific factors might be contributing to these results. Future studies are needed to identify the exact factors.

### Meta-analysis using MetaCyto identifies novel racial differences in immune cells

Our meta-analysis using the guided pipeline in MetaCyto revealed five cell types to be significantly different between the Asian and White. Asians have higher percentages of total CD4+ T cells and CD4+ central memory T cells, and lower percentages of natural killer (NK) cells, naïve CD8+ T cells and total CD8+ T cells (Fig. 5a). Among these findings, only the difference of total CD4+ T cells has been reported previously^33^. MetaCyto was able to identify this racial difference consistently across all cytometry panels (Fig. 5b). Combining the results from all panels allowed us to confirm the difference with high confidence (p = 1.2×10^−7^).

**Figure 5:**
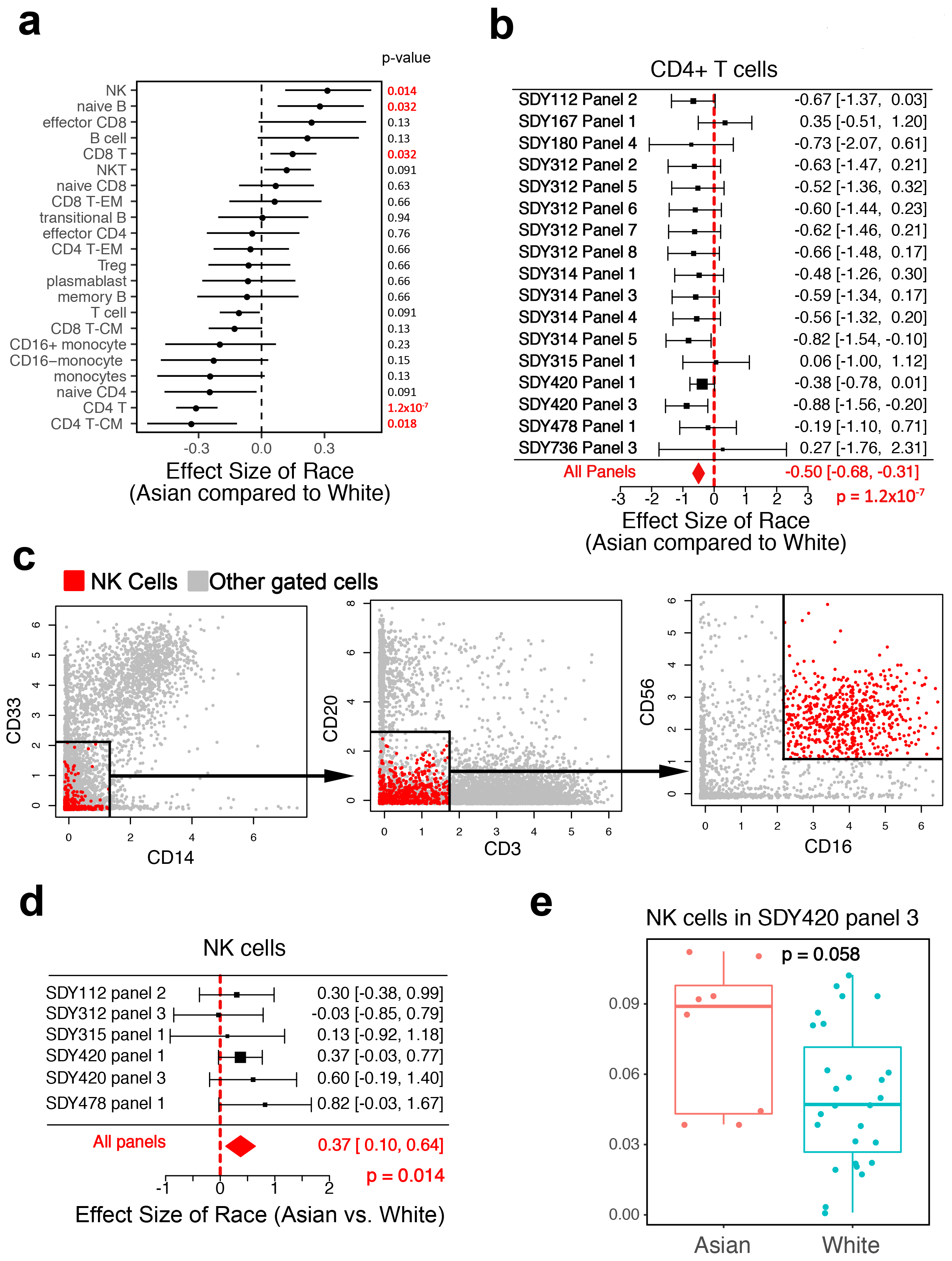
Meta-analysis of cytometry data using MetaCyto identifies multiple racial differences in immune cells. (**a**) A plot showing the effect size of race (Asian compare to White) on the proportion of 23 cell types in blood. Dots and whiskers represent the means and 95% confidence intervals. (**b**) A forest plot showing the effect size of race (Asian compared to White) on the proportion of CD4+ T cells in the blood. The effect sizes were estimated within each panel first, and are combined using a random effect model. (**c**) 2D plots visualizing the NK cells identified by MetaCyto. Red dots represent the cells identified by MetaCyto. Grey dots represent other cells in the gate. Data from SDY420 are shown as an example. (**d**) A forest plot showing the effect size of race on the proportion of NK cells in the blood. (**e**) The proportion of the NK cells in PBMC from Asian and White subjects in panel 3 of SDY420. The p values in **a, b and d** were calculated using random effect models, adjusted using Benjamini-Hochberg correction. p value in **e** is calculated from unpaired Mann-Whitney test without correction.

To confirm the four novel racial differences, we inspected the results from MetaCyto in detail. First, we visualized the identified cell populations in all studies, and confirmed that our results were not artifacts of automated gating (Fig. 5c and **Supplementary Fig. 4a**). We then tested if these racial differences were consistent across cytometry panels. Cochran’s Q test did not identify significant heterogeneity between cytometry panels (p value ranges from 0.27 to 0. 9). Visual inspection of the forest plots also confirmed that the results were consistent in most of the cytometry panels (Fig. 5d and **Supplementary Fig. 4b**).

Results from the unsupervised analysis identified multiple cell types, other than the 23 types used in the guided analysis, whose abundance were different between Asian and White (**Supplementary Table 6**). As one example, we found that the proportion of a subpopulation of CD8+ T cells, the CD3+ CD4− CD45RA+ CD8+ CD85J− cell population, is significantly higher in Asians than in Whites (**Supplementary Fig. 5**). A closer look at the forest plot revealed that the association between this population and race was not at a significant level in most studies taken independently. However, by combining the results from multiple studies, we were able to identify this association with high confidence (p=0.0049).

## DISCUSSION

With the collection of publically available cytometry studies rapidly growing, researchers can often identify multiple studies that were designed or can be re-purposed to answer a common research question. Meta-analysis of these studies allows researchers to answer the research question with a more robust conclusion and higher statistical power. Many cytometry studies that are publically available include hundreds of high dimensional cytometry data. Performing meta-analysis manually on these studies is not only time consuming, but also prone to human error and bias. In this study, we developed and demonstrated a computational tool called MetaCyto, which allows fully automated meta-analysis of both CyTOF and flow cytometry data.

When performing a meta-analysis of cytometry data, a big challenge lies in the identification of common cell subsets across heterogeneous cytometry studies. In MetaCyto, we implemented two complementary analysis pipelines to automate the cell identification process. The guided analysis pipeline is able to identify cell populations using user-defined cell definitions. For example, regulatory T cells can easily be identified using the definition “CD3+ CD4+ Foxp3+”. Such an approach allows researchers to incorporate their domain knowledge into the analysis, making the result more biologically relevant. In addition to the guided analysis pipeline, MetaCyto also allows researchers to identify cell populations in an un-supervised manor. Due to the high dimensionality of cytometry data, an exhaustive grid search will lead to an astronomical number of cell subsets. For example, if we divide each marker into positive and negative regions, 45 markers in a CyTOF experiment have 2^45^ combinations. To avoid such a situation, the unsupervised pipeline in MetaCyto first identifies cell clusters using a clustering method. Successful efforts were made by the community to develop efficient clustering methods for flow cytometry data analysis. We built MetaCyto to be fully compatible with existing clustering methods. MetaCyto is able to merge and transform the clusters from existing clustering algorithms in a biologically meaningful way, therefore improving result quality and enabling further meta-analysis of many studies.

Based on the test result, we recommend over-clustering the data first, followed by the merging of the clusters by MetaCyto. Such a strategy not only makes the method tuning free, but also is more computationally efficient than traditional auto-tuning methods, which require running the clustering algorithm multiple times with different parameters.

In MetaCyto, the distribution of each marker is bisected into positive and negative regions using a silhouette-scanning method. However, some markers may show tri-modal distributions. Although the silhouette-scanning method can easily be modified to divide the distribution into three regions (low-medium-high), only bisection is used in MetaCyto for the following reasons. First, it is known that multiple technical factors, such as auto-fluorescence, compensation, transformation and non-specific binding of antibodies, can lead to false tri-modal distributions^34,35^. In these cases, the low-medium-high regions do not represent distinct cell populations. Second, upon examining multiple cytometry studies, we found that although some markers (e.g. CD8, CD45RA, CD127) show tri-modal distributions in certain cytometry studies, they show bi-model distributions in other studies. Such inconsistency makes it difficult to reliably relate cell subsets across studies. Finally, our test result shows that bisection using silhouette scanning is able to identify the population that is truly positive for a marker even when the distribution is not bi-modal.

There are several potential limitations of the current study. In the unsupervised analysis pipeline of MetaCyto, although the merging step makes the clustering result more robust, it may eliminate some small cell populations of biological meaning. To overcome this limitation, researchers can use a more sensitive method, such as CITRUS^13^, to identify the cell subsets of interest from a single study. They can then craft cell definitions for those subsets and use the guided analysis pipeline of MetaCyto to perform meta-analysis across studies. Another limitation is that our meta-analysis only established correlations between cell populations and race. In-depth studies are needed to further validate our findings and to identify the genetic or environmental causes of these differences.

In summary, we developed MetaCyto, a computational tool that allows the automated meta-analysis of cytometry data. Applying MetaCyto to cytometry data from 10 human immunology studies allowed us to thoroughly characterize differences in the immune system between Asian and White populations. Other than the previously known differences in CD4+ T cell abundance, we identified novel cell populations whose abundance were significantly different between the two races, and demonstrated that the findings are consistent across multiple independent studies. Our findings can help us better understand the heterogeneity of the human immune system in the population. They also serve as a starting point for future studies to reveal the mechanisms behind racial discrepancies in immune-related diseases

## METHODS

### Data Aggregation

Flow cytometry data and CyTOF data from SDY112, SDY167^19^, SDY180^18^, SDY311, SDY312, SDY314, SDY315, SDY420^20^, SDY478 and SDY736^17^ were downloaded from ImmPort web portal. Only fcs files from pre-vaccination blood samples of healthy adults were included in the meta-analysis. Parameters, including antibodies and fluorescence or isotope labels, used in each fcs file were then identified using the *fcsInfoParser* function in MetaCyto. The fcs files were then organized into panels, which are defined as a collection of fcs files from the same study that have the same set of parameters.

Manual gating results for both FlowCAP WNV data (ID number FR-FCM-ZZY3) were downloaded from the FlowRepository link: community.cytobank.org/cytobank/experiments/4329.

All data sets were downloaded between September 1, 2016 and February 1, 2017.

### Data Pre-processing

Flow cytometry data from ImmPort were compensated for fluorescence spillovers using the compensation matrix supplied in each fcs file. All data from ImmPort were arcsinh transformed. For flow cytometry data, the formula f(x) = arcsinh (x/150) was used. For CyTOF data, the formula f(x) = arcsinh (x/8) was used. All transformation and compensation were done using the *preprocessing* or *preprocessing.batch* function in MetaCyto.

Cytometry data FlowCAP WNV was transformed and subset to only include protein markers. The pre-processing was doing using the same code provided by the Weber study^15^ : github.com/lmweber/cytometry-clustering-comparison

### Evaluating the performance of bisection algorithm

In **Supplementary Fig. 1**, Different Methods were tested to bisect the distribution of each marker.

Silhouette scanning method: The range of a marker was divided into 100 intervals using 99 breaks. The distribution was bisected at each break and the corresponding average silhouette^26^ was calculated. The break giving rise to the largest average silhouette was used as the cutoff for bisection.

K-means method: based on the values of a single marker, cells were clustered into 2 groups using k means clustering algorithm where k = 2. The cutoff value for bisection was the border between the 2 groups.

Hierarchical clustering method: based on the values of a single marker, cells were grouped into a Hierarchical tree. The tree was then cut into 2 groups at the top level. The cutoff value for bisection was the border between the 2 groups.

First valley method: The distribution of each marker was smoothed using the *smooth.spline* function. The peaks in the distribution were identified using the *.getPeaks* function in flowDensity package^27^. The lowest points between peaks were defined as valleys. The valley with the smallest marker value was used as the cutoff for bisection.

Last valley method: The valley with the largest marker value was used as the cutoff for bisection.

Median valley method: The valley closest to the median of the marker value was used as the cutoff for bisection.

Mean method: The mean of the marker distribution was used as the cutoff.

Median method: the median of the marker value was used as the cutoff

Middle method: the mean of the max and min of the marker values were used as the cutoff.

After markers in SDY420 data were bisected, cells fulfilling the requirement of each cell definition (listed in **Supplementary Table 1**) were identified. For example, for cell definition “CD3+ CD8+ CD4−”, cells falling into the CD3+ region were identified. Similarly, cells falling into CD8+ and CD4− regions were identified. The intersection of the 3 sets of cells was the cells corresponding to the cell definition “CD3+ CD8+ CD4−”. The proportion of cells corresponding to each cell definition was calculated and compared to the proportion provided by the author. The Spearman correlation was used as a measurement of the bisection algorithm.

### Identifying cell subsets with the guided analysis pipeline in MetaCyto

Cell definitions were created based on the gating strategies provided by authors of SDY 420 and SDY478 from ImmPort database or based on the cell definition from the Human ImmunoPhenotyping Consortium^29^. The cell definitions are available in the **supplementary table 1, 2** and **4**.

To identify the corresponding cell subsets, Silhouette scanning was used to bisect the distribution of cell markers into positive and negative regions. Cells fulfilling the cell definitions were then identified. For example, the CD3+ CD4+ CD8− (CD4+ T cells) cell subset corresponds to the cells that fall into the CD3+ region, CD4+ region and CD8− region concurrently. The proportion of each cell subset in blood was calculated by dividing the number of cells in the subset by the total number of cells in the blood. The procedure is performed using the *searchCluster* or *searchClster.batch* function in the Metacyto package.

### Identifying cell subsets with the un-supervised analysis pipeline in MetaCyto

FlowSOM^11^ or FlowMeans^12^ were used to identify cell clusters in the cytometry data. Silhouette scanning was used to identify a threshold that bisects the distribution of cell markers into positive and negative regions. To label the identified cell clusters, the marker levels in each cluster were compared with the bisection threshold. If the marker levels of 95% of cells in the cluster are above or below the threshold, the cluster will be labeled as positive or negative for the marker, respectively. Otherwise, the cluster will not be labeled for the marker. If a marker is positive or negative in 95% of all cells, the marker is not used to label any clusters. The procedure is performed using the *labelCluster* function in MetaCyto.

The generated labels were then used to identify the corresponding cell subsets, in a fashion similar to the guided analysis pipeline. Notice that such a process is equivalent of merging cell clusters that have the same labels into a hyper-rectangle shaped cluster. The proportion of each cell subset in blood was calculated by dividing the number of cells in the subset by the total number of cells in the blood. The procedure was performed using the *searchCluster* or *searchClster.batch* function in the Metacyto package.

### Comparing guided analysis pipeline in MetaCyto with flowDensity and ACDC

Cell definitions were created based on the gating strategies provided by authors of. The cell definitions are available in the **supplementary table 2**.

For MetaCyto, the proportion of each cell subset in blood was estimated by the guided analysis pipeline described above.

For flowDensity, the *flowDensity* function was used to identify the cell subsets corresponding to the cell definitions. The proportion of each cell subset in blood was calculated by dividing the number of cells in the subset by the total number of cells in the blood.

For ACDC, the cell definitions were first turned into a cell type-marker table with entries of 1, −1, and 0 representing a marker is present, absent, and ignored in a cell type respectively. The landmark points of each cell type were generated by clustering samples within each partition defined by the cell type-marker table. Landmark points were used to classify samples with the semi-supervised classification algorithm. Only the bottom level cell types (basophils, NK cells, naive B cells, memory B cells, transitional B cells, plasmablasts, effector memory CD4+ T cells, effector CD4+ T cells, central memory CD4+ T cells, naive CD4+ T cells, Tregs, NKT cells, effector memory CD8+ T cells, effector CD8+ T cells, central memory CD8+ T cells, naive CD8+ T cells, gamma-delta T cells and monocytes) were estimated by the ACDC because it is designed to detect mutually exclusive cell types. Their parent populations, such as total T cells, were excluded.

All three methods received the same types of input: preprocessed CyTOF data and the cell definitions. No additional information or human input was provided to help identifying cell subsets. The Spearman correlation coefficient between author’s result and MetaCyto or flowDensity or ACDC result was calculated to measure the performance.

### Comparing un-supervised analysis pipeline in MetaCyto with FlowSOM and FlowMeans

FlowSOM^11^ and FlowMeans^12^ used to cluster cytometry data using different K (number of clusters) values. The resulting clusters from FlowSOM and FlowMeans were labeled and merged using the same procedure described in the un-supervised pipeline.

The F measure was used to measure the performance of clustering methods with or with out the merging step. The F measure was calculated as described in the FlowCAP study^36^. Briefly, for each cell population in the manual gating result and each cell population in the auto-gating result, a 2 × 2 contingency table was calculated containing the false positive (FP), true positive (TP), false negative (FN) and true negative (TN). The recall (Re) was calculated as TP/(TP + FN), the precision (Pr) was calculated a TP/(TP + FP). The F measure was calculated as *F* = (2 × Pr × Re)/(Pr + Re). For each population in manual gating result, the best F measure and its corresponding recall and precision were used as the F measure of the population. The overall F measure, Recall and Precision were the average of F measure, Recall and Precision of all manual gated populations, weighted by the size of each manual population.

### Estimating the proportions of cell subsets using data from Carr study

The cell gating results from Carr study, in the form of proportions of parents gate, are available as Supplementary Data Set 1 of the Carr study pulication^30^. The proportion of a cell subset within blood is calculated by multiplying the proportions of parents in all gating steps together. For example, the proportion of CD4 T cell in blood is calculated by multiplying the proportion of total T cells in blood by the proportion of CD4 T cells in total T cells.

### Statistical Analysis

2-level hieratical regression models were used in the meta-analysis of the 10 human immunology studies from ImmPort: the proportion of cell subsets was regressed against age, gender and race (Y ~ age + gender + race) in each cytometry panel. The effect size was defined as the regression coefficient divided by the standard deviation of Y. The overall effect size from all cytometry panels was estimated using a random effect model. For data from the Carr study, the proportion of a cell population was regressed against age and gender. Race information was missing in the data, therefore was omitted in the regression. All statistical analysis was performed using the *metaAnalysis* function in MetaCtyo. The p-value was adjusted using the Benjamini-Hochberg^37^ correction.

To test the heterogeneity in Meta-analysis, Cochran’s Q test was performed using the *cochran.Q* function in the *Mada* package.

In Fig. 4 a **and b**, Pearson correlations are calculated and tested against the null hypothesis (correlation equals zero) using the *cor.test* function in R.

In Fig. 5e **and Supplementary Fig. 5b**, Shapiro-Wilk test was performed to check the normality assumption using the *shapiro.test* function in R. F test was performed to check the equal variance assumption using the *var.test* function in R. A two-sided unpaired Mann-Whitney test is performed to test the difference between two groups using the *wilcox.test* function in R.

### Code availability

The MetaCyto R package is available on GitHub: github.com/hzc363/MetaCyto

The source code to reproduce results in Figure 2-5 is available on GitHub: github.com/hzc363/MetaCyto_Paper_Code

## SUPPLEMENTARY INFORMATION

Supplementary Figure 1: Silhouette scanning bisects the distribution of each marker in a biologically meaningful way. (**a-c**) Examples illustrating how silhouette scanning bisects markers with bimodal distribution (**a**), tri-modal distribution (**b**) and a distribution where the positive population does not form a separate peak (**c**). The range of a marker is divided into 100 intervals using 99 breaks. The distribution is bisected at each break and the corresponding average silhouette is calculated. The break that gives rise to the largest average silhouette is used as the cutoff for bisection. Grey histogram shows the distribution of the markers. Blue dots show the average silhouette at each break. Red line shows the cutoff that maximizes the average silhouette. Black arrows show the position of 3 peaks in (b). (**d-e**) Using different bisection algorithms, each marker in CyTOF data from SDY420 are bisected into positive and negative regions. 24 cell types were identified using the guided analysis pipeline as described in Fig. 1c. The proportion of each cell type in each sample is calculated and compared with manual gating result. (**d**) The Spearman correlation between the estimated proportion and author’s proportion are used to measure the performance of each bisection algorithm. (**e**) Scatter plots showing the result generated by using silhouette scanning, k-means clustering and mean as the bisection algorithm. Each dot represents the proportion of a cell type in a sample. Each color represents a cell type. See online methods for a detailed description of the 9 bisection methods tested.

Supplementary Figure 2: MetaCyto accurately identifies cell populations (**a-b**) Scatter plots showing the comparison between proportions of cell types estimated by the guided analysis pipeline in MetaCyto (**a**) or ACDC (**b**) and proportions provided by the authors of SDY478. Each dot represents the proportion of a cell type in a sample. Each color represents a cell type. (**c-d**) FlowMeans is used to cluster FlowCAP WNV data with K ranging from 10 to 90 with or without MetaCyto. F measure (**c**) and the number of clusters (**d**) are showed in the bar plots. (**e-f**) FlowSOM is used to cluster FlowCAP ND data with K ranging from 10 to 90 with or without MetaCyto. F measure (**e**) and the number of clusters (**f**) are showed in the bar plots.

Supplementary Figure 3: Demographics of subjects included in the meta-analysis of 10 human immunology studies. Bar graphs show the distribution of gender (**a**), race (**b**) and age (**c**).

Supplementary Figure 4: Racial differences identified by MetaCyto are consistent across cytometry panels. (**a**) Representative 2D plots visualizing the cell subsets (Naïve B cells, CD8+ T cells and CD4+ central memory T cells) identified by MetaCyto. Red dots represent the cells identified by MetaCyto. Grey dots represent other cells in the gate. Data from SDY420 are shown as examples. (**b**) Forest plots showing the effect size of race (Asian compare to White) estimated in each cytometry panel for Naïve B cells, CD8+ T cells and CD4+ central memory T cells. The effect sizes were estimated within each panel first, and were combined using a random effect model.

Supplementary Figure 5: An example of racial differences identified by the unsupervised pipeline. (**a**) Forest plot showing the effect size of race (Asian vs. White) on the proportion of a novel cell subset (CD3+CD4-CD45RA+CD8+CD85J-) identified by the unsupervised analysis pipeline. (**b**) The proportion of the CD3+CD4-CD45RA+CD8+CD85J− cell subset in PBMC from Asian and White subjects in panel 2 of SDY312. The p value in a is calculated using a random effect model, adjusted using Benjamini-Hochberg correction. p value in **b** is calculated from unpaired Mann-Whitney test without correction. See **Supplementary Table 6** for a complete list of differences in immune cells between races identified by the unsupervised pipeline.

Supplementary Table 1: A list of cell definitions used to identify the 24 cell populations in cytometry data (SDY420) from ImmPort. The cell definitions are created based on the author’s gating strategy provided in SDY420.

Supplementary Table 2: A list of cell definitions used to identify the 88 cell populations in cytometry data (SDY478) from ImmPort. The cell definitions are created based on the author’s gating strategy provided in SDY478.

Supplementary Table 3: A summary of 10 studies included in the meta-analysis.

Supplementary Table 4: A list of cell definitions used to identify the 23 cell populations in all 10 studies included in the meta-analysis.

Supplementary Table 5: A table reporting the Pearson correlation between the results from leave-one-out analysis and the results from the full meta-analysis

Supplementary Table 6: A table summarizing the effect size of race to the proportion of cell subsets identified by the unsupervised analysis pipeline when comparing cytometry data of blood from Asian and White subjects. The effect sizes are estimated from the meta-analysis of 10 human immunology studies.

## ACKNOWLEDGMENTS

We thank Marina Sirota, Dvir Aran, Henry Schaefer, Elizabeth Thomson, Kelly Zalocusky and Matthew Kan for helpful discussion.

Research reported in this publication was supported by the National Institute of Allergy and Infectious Diseases (Bioinformatics Support Contract HHSN272201200028C). The content is solely the responsibility of the authors and does not necessarily represent the official views of the National Institutes of Health.

## AUTHOR CONTRIBUTIONS

Z.H., A.J.B. and S.B. conceived the study and developed the method. Z.H. and C.J. designed the overall structure of MetaCyto and performed the meta-analysis of cytometry data from ImmPort. J.H., M.S., P.F.G., S.A., P.D., C.T., J.W., G.N. gave valuable input and suggestions for analysis. H.L., B.A.K. and J.T.D. evaluated the performance of ACDC. H.Z. and C.J. wrote the manuscript and made figures with input from co-authors.

## REFERENCES

1. Sutton, A. J., Abrams, K. R., Jones, D. R. & Sheldon, T. A. Methods for Meta-analysis in Medical Research Contents Preface Acknowledgements Part A: Meta-Analysis Methodology: The Basics.

2. Wirapati, P. et al. Meta-analysis of gene expression profiles in breast cancer: toward a unified understanding of breast cancer subtyping and prognosis signatures. Breast Cancer Res. 10, R65 (2008).

3. Kodama, K. et al. Expression-based genome-wide association study links the receptor CD44 in adipose tissue with type 2 diabetes. Proc. Natl. Acad. Sci. U. S. A. 109, 7049–54 (2012).

4. Boulé, N. G., Haddad, E., Kenny, G. P., Wells, G. A. & Sigal, R. J. Effects of Exercise on Glycemic Control and Body Mass in Type 2 Diabetes Mellitus. JAMA 286, 1218 (2001).

5. Brown, S. D. et al. Neo-antigens predicted by tumor genome meta-analysis correlate with increased patient survival. Genome Res. 24, 743–50 (2014).

6. Barrett, J. C. et al. Genome-wide association study and meta-analysis find that over 40 loci affect risk of type 1 diabetes. Nat. Genet. 41, 703–707 (2009).

7. Perfetto, S. P., Chattopadhyay, P. K. & Roederer, M. Innovation: Seventeen-colour flow cytometry: unravelling the immune system. Nat. Rev. Immunol. 4, 648–655 (2004).

8. Shapiro, H. M. Multistation multiparameter flow cytometry: A critical review and rationale. Cytometry 3, 227–243 (1983).

9. Bandura, D. R. et al. Mass Cytometry: Technique for Real Time Single Cell Multitarget Immunoassay Based on Inductively Coupled Plasma Time-of-Flight Mass Spectrometry. Anal. Chem. 81, 6813–6822 (2009).

10. Bhattacharya, S. et al. ImmPort: disseminating data to the public for the future of immunology. Immunol. Res. 58, 234–239

11. Van Gassen, S. et al. FlowSOM: Using self-organizing maps for visualization and interpretation of cytometry data. Cytom. Part A 87, 636–645 (2015).

12. Aghaeepour, N., Nikolic, R., Hoos, H. H. & Brinkman, R. R. Rapid cell population identification in flow cytometry data. Cytom. Part A 79A, 6–13 (2011).

13. Bruggner, R. V., Bodenmiller, B., Dill, D. L., Tibshirani, R. J. & Nolan, G. P. Automated identification of stratifying signatures in cellular subpopulations. Proc. Natl. Acad. Sci. 111, E2770–E2777 (2014).

14. Aghaeepour, N. et al. Critical assessment of automated flow cytometry data analysis techniques. Nat. Methods 10, 228–238 (2013).

15. Weber, L. M. & Robinson, M. D. Comparison of Clustering Methods for High-Dimensional Single-Cell Flow and Mass Cytometry Data. bioRxiv 47613 (2016). doi:10.1101/047613

16. Qiu, P. et al. Extracting a cellular hierarchy from high-dimensional cytometry data with SPADE. Nat. Biotechnol. 29, 886–891 (2011).

17. Wertheimer, A. M. et al. Aging and Cytomegalovirus Infection Differentially and Jointly Affect Distinct Circulating T Cell Subsets in Humans. J. Immunol. 192, 2143–2155 (2014).

18. Obermoser, G. et al. Systems Scale Interactive Exploration Reveals Quantitative and Qualitative Differences in Response to Influenza and Pneumococcal Vaccines. Immunity 38, 831–844 (2013).

19. Ledgerwood, J. E. et al. Influenza virus h5 DNA vaccination is immunogenic by intramuscular and intradermal routes in humans. Clin. Vaccine Immunol. 19, 1792–7 (2012).

20. Whiting, C. C. et al. Large-Scale and Comprehensive Immune Profiling and Functional Analysis of Normal Human Aging. PLoS One 10, e0133627 (2015).

21. Achhra, A. C. et al. Difference in absolute CD4+ count according to CD4 percentage between Asian and Caucasian HIV-infected patients. J. AIDS Clin. Res. 1, 1–4 (2010).

22. Petri, M. Epidemiology of systemic lupus erythematosus. Best Pract. Res. Clin. Rheumatol. 16, 847–58 (2002).

23. Golden-Mason, L., Klarquist, J., Wahed, A. S. & Rosen, H. R. Cutting edge: programmed death-1 expression is increased on immunocytes in chronic hepatitis C virus and predicts failure of response to antiviral therapy: race-dependent differences. J. Immunol. 180, 3637–41 (2008).

24. Li, Y. et al. Inter-individual variability and genetic influences on cytokine responses to bacteria and fungi. Nat. Med. 22, 952–60 (2016).

25. Brodin, P. et al. Variation in the Human Immune System Is Largely Driven by Non-Heritable Influences. Cell 160, 37–47 (2015).

26. Rousseeuw, P. J. Silhouettes: A graphical aid to the interpretation and validation of cluster analysis. J. Comput. Appl. Math. 20, 53–65 (1987).

27. Malek, M. et al. flowDensity: reproducing manual gating of flow cytometry data by automated density-based cell population identification. Bioinformatics 31, 606–607 (2015).

28. Lee, H.-C., Kosoy, R., Becker, C. E., Dudley, J. T. & Kidd, B. A. Automated cell type discovery and classification through knowledge transfer. Bioinformatics 11, 1822–1833 (2017).

29. Finak, G. et al. Standardizing Flow Cytometry Immunophenotyping Analysis from the Human ImmunoPhenotyping Consortium. Sci. Rep. 6, 20686 (2016).

30. Carr, E. J. et al. The cellular composition of the human immune system is shaped by age and cohabitation. Nat. Immunol. 17, 461–468 (2016).

31. Yan, J. et al. The effect of ageing on human lymphocyte subsets: comparison of males and females.

32. Uppal, S. S., Verma, S. & Dhot, P. S. Normal values of CD4 and CD8 lymphocyte subsets in healthy indian adults and the effects of sex, age, ethnicity, and smoking. Cytometry 52B, 32–36 (2003).

33. Howard, R. R., Fasano, C. S., Frey, L. & Miller, C. H. Reference intervals of CD3, CD4, CD8, CD4/CD8, and absolute CD4 values in asian and non-asian populations. Cytometry 26, 231–232 (1996).

34. Ray, S. & Pyne, S. A Computational Framework to Emulate the Human Perspective in Flow Cytometric Data Analysis. PLoS One 7, (2012).

35. Morice, W. G. et al. Flow Cytometric Assessment of TCR-V b Expression in the Evaluation of Peripheral Blood Involvement by T-Cell Lymphoproliferative Disorders: A Comparison With Conventional T-Cell Immunophenotyping and Molecular Genetic Techniques. Am. J. Clin. Pathol. 121, 373–383 (2004).

36. Aghaeepour, N. et al. Critical assessment of automated flow cytometry data analysis techniques. Nat. Methods 10, 228–238 (2013).

37. Author, T., Benjamini, Y., Hochberg, Y. & Benjaminit, Y. Controlling the False Discovery Rate: A Practical and Powerful Approach to Multiple Controlling the False Discovery Rate: a Practical and Powerful Approach to Multiple Testing. Source J. R. Stat. Soc. Ser. B J. R. Stat. Soc. Ser. BMethodological) J. R. Stat. Soc. B 57, 289–300 (1995).

